# *retro*-Tango enables versatile retrograde circuit tracing in *Drosophila*

**DOI:** 10.1101/2022.11.24.517859

**Authors:** Altar Sorkaç, Rareș A Moșneanu, Anthony M Crown, Doruk Savaş, Angel M Okoro, Mustafa Talay, Gilad Barnea

## Abstract

Transsynaptic tracing methods are crucial tools in studying neural circuits. Although a couple of anterograde tracing methods and a targeted retrograde tool have been developed in *Drosophila melanogaster*, there is still need for an unbiased, user-friendly, and flexible retrograde tracing system. Here we describe *retro*-Tango, a method for transsynaptic, retrograde circuit tracing and manipulation in *Drosophila*. In this genetically encoded system, a ligand-receptor interaction at the synapse triggers an intracellular signaling cascade that results in reporter gene expression in presynaptic neurons. Importantly, panneuronal expression of the elements of the cascade renders this method versatile, enabling its use not only to test hypotheses but also to generate them. We validate *retro*-Tango in various circuits and benchmark it by comparing our findings with the electron microscopy reconstruction of the *Drosophila* hemibrain. Our experiments establish *retro*-Tango as a key method for circuit tracing in neuroscience research.

## INTRODUCTION

The Turkish poet Nazım Hikmet wrote:

*To live, like a tree one and free*

*And like a forest, sisterly* (Hikmet, 2002).

This also holds true to the function of the nervous system. Like forests, neural circuits have evolved as congruous networks of individual units: neurons. These networks integrate external stimuli with the internal state of the animal and generate the proper behavioral responses to the changing environment. Therefore, understanding the individual neuron is invaluable for deciphering animal behavior; yet the study of circuits is an indispensable complement to it.

The study of neural circuits encompasses a variety of approaches of which the analysis of connectivity between neurons is fundamental. In this respect, the complete electron microscopy (EM) reconstruction of the *Caenorhabditis elegans* nervous system in the 1980s (White et al., 1986) and the ongoing efforts to complete the *Drosophila melanogaster* connectome (Bates, Schlegel, et al., 2020; Eichler et al., 2017; Engert et al., 2022; Fushiki et al., 2016; Horne et al., 2018; Hulse et al., 2021; Marin et al., 2020; Ohyama et al., 2015; Scheffer et al., 2020; Takemura, Aso, et al., 2017; Takemura, Nern, et al., 2017; Zheng et al., 2018) provide the gold standard for the analysis of neural circuits. These endeavors open new paths for the study of nervous systems. However, like all methods, they come with their own shortcomings.

The EM reconstruction of the *C. elegans* nervous system was originally performed with a single hermaphrodite reared at specific laboratory conditions. Further, it was not until 30 years later that the nervous system of a second animal, a male, was reconstructed (Cook et al., 2019). As to *D. melanogaster*, the brain of a single female is still being reconstructed. These time-consuming and labor-intensive aspects of EM reconstructions preclude the study of individual differences that might arise from variances such as sex, genetics, epigenetics, rearing conditions, and past experiences. Hence, transsynaptic tracing techniques remain valuable even in the age of EM connectomics.

In *D. melanogaster*, techniques such as photoactivatable GFP (PA-GFP) (Datta et al., 2008; Patterson & Lippincott-Schwartz, 2002) and GFP-reconstitution across synaptic partners (GRASP) (Fan et al., 2013; Feinberg et al., 2008; Gordon & Scott, 2009; Macpherson et al., 2015; Shearin et al., 2018) have been instrumental in studying neural circuits and connectivity. Recently, two methods, *trans-*Tango (Talay et al., 2017) and TRACT (Huang et al., 2017), were developed for anterograde transsynaptic tracing. In addition, a retrograde transsynaptic tracing method, termed BAcTrace, was devised (Cachero et al., 2020). All three techniques differ from the aforementioned PA-GFP and GRASP in that they provide genetic access to synaptic partners of a set of neurons, enabling their use in not only tracing but also monitoring and manipulation of neural circuits (Snell et al., 2022). Furthermore, *trans-*Tango and TRACT do not necessitate hypotheses prior to experimentation, since all neurons are capable of revealing the postsynaptic signal should the cascades be triggered by their presynaptic partners. In contrast, BAcTrace, by design, relies on the expression of the presynaptic components of the cascade solely in candidate neurons. Therefore, it requires a hypothesis to be tested, rendering this technique inherently biased. In addition, BAcTrace experiments are constrained by the availability of drivers in candidate neurons because the presynaptic components are expressed under a LexA driver. Hence, there is still a need for a versatile retrograde tracing method that can be used as a hypothesis tester, and, more importantly, as a hypothesis generator.

To fill this gap, here we present *retro*-Tango, a retrograde version of *trans-*Tango, as a user-friendly, versatile retrograde transsynaptic tracing technique for use in *D. melanogaster*. Like *trans-*Tango, *retro*-Tango functions through a signaling cascade initiated by a ligand-receptor interaction at the synapse and resulting in reporter expression in synaptic partners. To target the reporter expression to presynaptic neurons, we devised a ligand tethered to a protein that localizes to dendrites in the starter neurons.

In order to benchmark the system, we used it in various known circuits. First, we revealed the presynaptic partners of the giant fiber from the escape circuit and compared our results to the EM reconstruction. Second, to demonstrate the versatility of *retro*-Tango, we implemented it in the central complex. Third, we tested the specificity of the system by using it in a sexually dimorphic circuit where the presynaptic partners of a set of neurons differ between males and females. Lastly, we used *retro*-Tango in the sex peptide circuit and in the olfactory system where we traced connections from the central nervous system (CNS) to the periphery and vice versa. Our study establishes *retro*-Tango as a prime method for neuroscience research in fruit flies.

## RESULTS

### Design of *retro*-Tango

*retro*-Tango is the retrograde counterpart of the transsynaptic tracing technique *trans-* Tango (Talay et al., 2017), and both are based on the Tango assay for G-protein coupled receptors (GPCRs) (Barnea et al., 2008). In the Tango assay, activation of a GPCR by its ligand is monitored via a signaling cascade that eventually results in reporter gene expression. This signaling cascade comprises two fusion proteins. The first is a GPCR tethered to a transcriptional activator via a cleavage site recognized by the tobacco etch virus N1a protease (TEV). The second is the human β-arrestin2 protein fused to TEV (Arr::TEV). A third component is a reporter gene under control of the transcriptional activator. Upon binding of the ligand to the receptor, arrestin is recruited to the activated receptor bringing TEV in close proximity to its recognition site. TEV-mediated cleavage then releases the transcriptional activator that in turn translocates to the nucleus to initiate transcription of the reporter gene. These components are conserved in both transsynaptic tracing techniques, *trans-*Tango (Talay et al., 2017) and *retro*-Tango. The novelty in both methods is in the tethering of the ligand to a transmembrane protein to localize it to pre- (*trans-*Tango), or post- (*retro*-Tango) synaptic sites. In this manner, the ligand activates its receptor only across the synaptic cleft and initiates the signaling cascade in synaptic partners. In both methods, the human glucagon (GCG) and the human glucagon receptor (GCGR) are used as the ligand-receptor pair, and the GCGR is tethered to the transcriptional activator QF (GCGR::TEVcs::QF) (Figure 1a).

**Figure 1.**
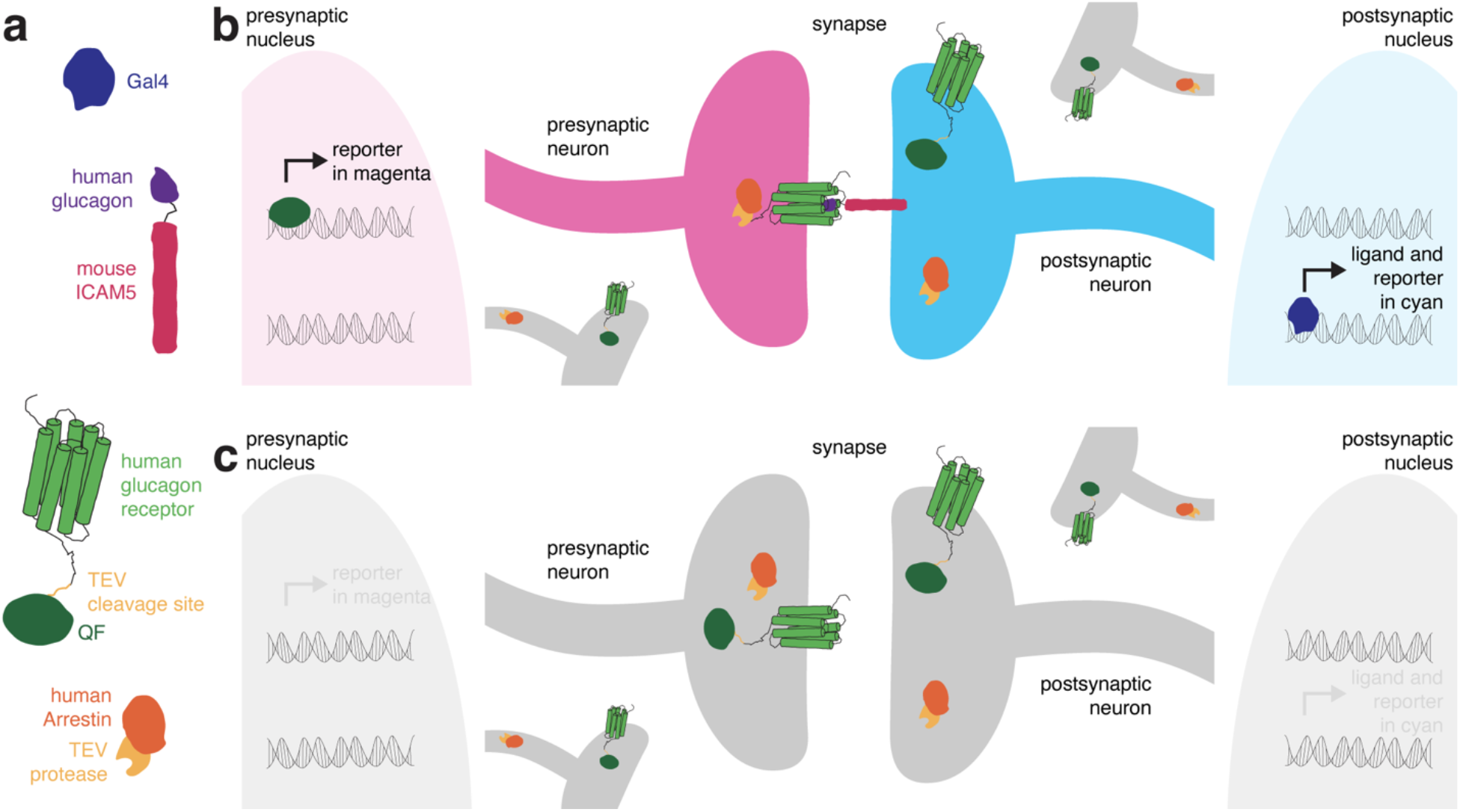
The design of *retro*-Tango. **(a)** The components of *retro*-Tango. **(b)** In *retro*-Tango, all neurons express two of the components of the signaling cascade: human glucagon receptor::TEV cleavage site::QF and human β-arrestin2::TEV protease. They also carry the gene encoding the presynaptic mtdTomato reporter (magenta) under the control of QF. Therefore, all neurons are capable of expressing the reporter. In starter neurons expressing Gal4, the ligand (human glucagon::mouse ICAM5) is expressed along with the GFP reporter (cyan) marking the postsynaptic starter neurons. The mICAM5 fusion localizes the ligand to the postsynaptic sites such that the ligand activates its receptor only across the synapse. Upon activation of the receptor in the presynaptic neuron, the Arrestin-TEV fusion is recruited. TEV-mediated proteolytic cleavage then releases the transcription factor QF from the receptor. QF in turn translocates to the nucleus and initiates transcription of the presynaptic magenta reporter. In neurons that are not presynaptic to the starter neurons, the reporter is not expressed. **(c)** In the absence of a Gal4 driver, the ligand is not expressed, and the signaling cascade is not triggered, resulting in no expression of the reporters.

In *retro*-Tango the targeting of glucagon to postsynaptic sites is achieved via the mouse intercellular adhesion molecule ICAM5 (Figure 1a). In *Drosophila* neurons, this protein is present at low levels in cell bodies and mainly localizes to the dendrites but not the axons, enabling its use as a dendritic marker (Nicolai et al., 2010). The ligand and the postsynaptic reporter farnesylated GFP are stoichiometrically expressed under the control of the Gal4/UAS system using the self-cleaving P2A peptide (Daniels et al., 2014) (Figure 1–figure supplement 1). In this manner, the presence of the ligand is coupled with the GFP signal, eliminating any discrepancy that might arise from differentially expressing them from two separate genomic sites. Both the GCGR::TEVcs::QF and the Arr::TEV fusion proteins are expressed panneuronally, and the expression of the presynaptic reporter mtdTomato is controlled by the QF/QUAS binary system (Potter et al., 2010) (Figure 1–figure supplement 1). In postsynaptic starter cells, Gal4 drives the expression of both GFP and the ligand (Figure 1b). The interaction of the ligand with its receptor on the presynaptic partners triggers the *retro*-Tango cascade that culminates in mtdTomato expression in these neurons. By contrast, the ligand is not expressed in the absence of a Gal4 driver. Therefore, the cascade is not triggered, and no presynaptic signal is observed (Figure 1c). Since the presynaptic components of the pathway are expressed panneuronally, all neurons have the capacity to reveal the presynaptic signal when the ligand is expressed by their postsynaptic partners. Thus, the design of *retro*-Tango is not inherently biased.

### Validation of *retro*-Tango

For the initial validation of *retro*-Tango we chose the giant fibers (GFs) of the escape circuit. The GFs are descending command interneurons that respond to neural pathways sensing looming stimuli, such as from a predator. They then relay this information to downstream neurons for the fly to initiate the take-off response (Fotowat et al., 2009; von Reyn et al., 2014). The GFs receive direct input from two types of visual projection neurons: lobula columnar type 4 (LC4) (von Reyn et al., 2017) and lobula plate/lobula columnar type 2 (LPLC2) (Ache et al., 2019). They then integrate this information and convey it to the tergotrochanteral motor neurons (TTMns) and the peripherally synapsing interneurons (PSIs) in the ventral nerve cord (VNC). The GFs form chemical and electrical synapses with both of these types of neurons (Allen et al., 2006). All of these neurons are easily identifiable based on their morphology in the optic lobes or the VNC, rendering the GF system attractive for validating *retro*-Tango. In addition, there is a specific driver line that expresses only in the GFs (von Reyn et al., 2014). Further, the GFs are clearly annotated in the EM reconstruction of the hemibrain (Zheng et al., 2018), allowing for the comparison of the *retro*-Tango results with the annotated connectome.

When we initiated *retro*-Tango from the GFs in adult males, we observed strong presynaptic signal in cells with dense arborizations in the brain and sparse processes in the VNC (Figure 2a). Upon close examination, we noticed few cell bodies in the VNC, suggesting that the VNC signal originates mostly from descending neurons with somata in the brain. As expected, we did not observe *retro*-Tango signal in the TTMns and PSIs, known postsynaptic partners of the GFs. Importantly, we could identify neurons in the optic lobes with the characteristic dendritic arborizations of the LC4s and the LPLC2s, established presynaptic partners of the GFs. It is noteworthy that we observed sporadic asymmetrical signal in the postsynaptic starter neurons, a phenomenon we notice when we use some split-Gal4 drivers. Likewise, we observe asymmetry in the *retro*-Tango signal in the presynaptic neurons. The stronger signals in the postsynaptic and the presynaptic neurons are in the same hemisphere, likely reflecting higher ligand expression in the starter neurons. Such differences in signal intensity may lead to qualitative differences in presynaptic neurons revealed in each hemisphere. For example, the LC4 neurons (marked by the arrow) are visible only in one hemisphere (Figure 2a). Nonetheless, we conclude that *retro*-Tango yields strong signal and labels the expected presynaptic partners of the GFs. Further, it does not exhibit false positive signal in the postsynaptic targets of the GFs. These results indicate that *retro*-Tango is indeed selective to the retrograde direction.

**Figure 2.**
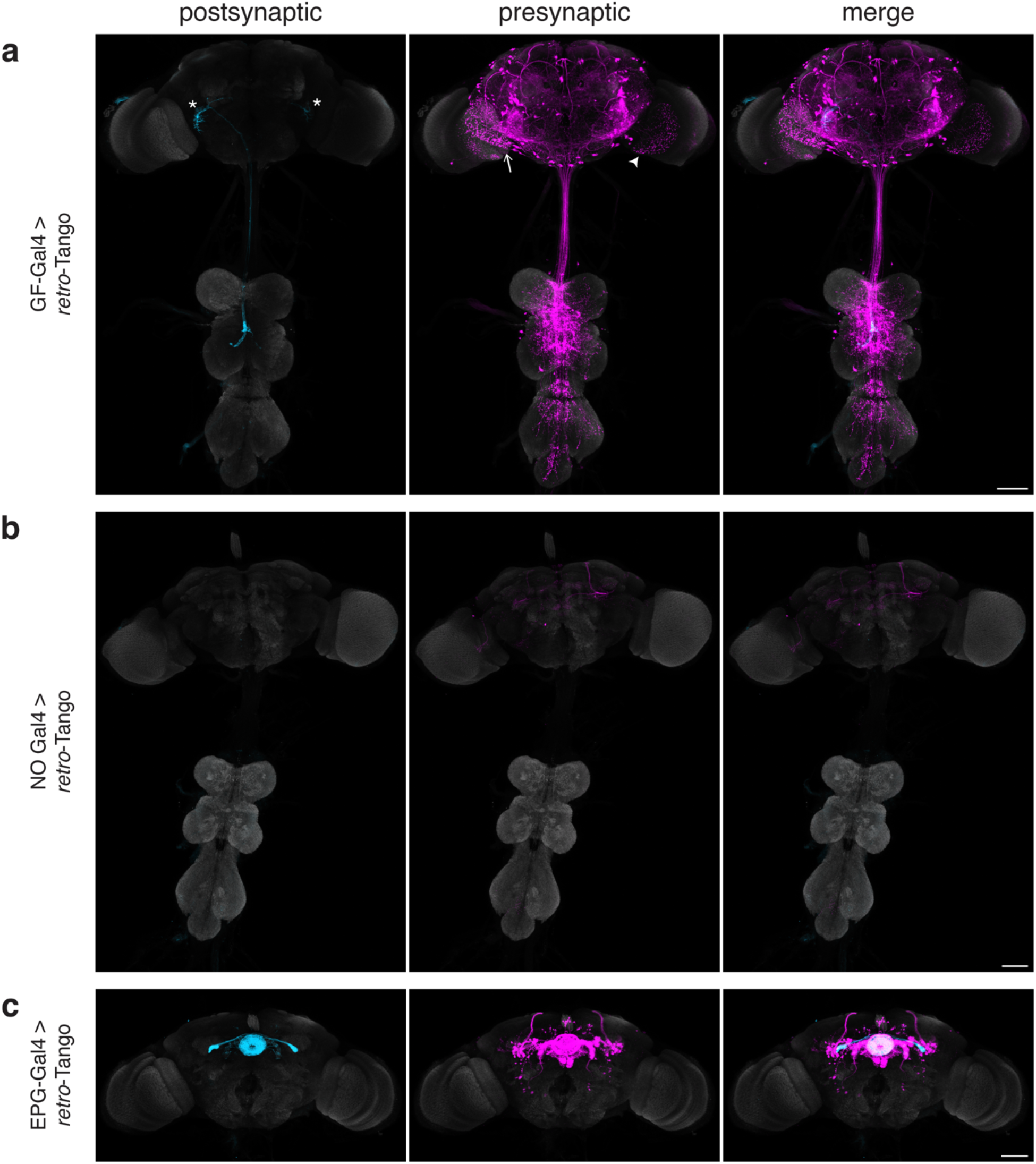
Implementation of *retro*-Tango in the giant fiber and central complex circuits. **(a)** Initiating *retro*-Tango from the GFs (asterisks mark the cell bodies) results in presynaptic signal in the brain and VNC. Both LC4 (arrow) and LPLC2 (arrowhead) neurons, known presynaptic partners of GFs, are identified by *retro*-Tango. Note the asymmetry between hemispheres in the signal in the postsynaptic starter neurons and their corresponding presynaptic partners. **(b)** *retro*-Tango exhibits little background noise in the absence of a Gal4 driver. Background is observed in the mushroom bodies, in the central complex, and in a few neurons in the VNC. **(c)** Ligand expression in EPG neurons of the central complex leads to *retro*-Tango signal in their known presynaptic partners: PEN, PFR and Δ7 neurons. The signal in these neurons can be easily discerned from the background noise. Postsynaptic GFP (cyan), presynaptic mtdTomato (magenta) and neuropil (grey). Scale bars, 50μm.

It is noteworthy that we do not observe strong background noise with *retro*-Tango in the absence of a Gal4 driver where the ligand is not expressed (Figure 2b). There is, however, faint background noise in some of the Kenyon cells of the mushroom body as well as in the fan-shaped body and noduli of the central complex. In addition, we occasionally observe sporadic noise in a few neurons in the VNC. This background noise might be due to leaky expression of the ligand, albeit in low levels as reflected by the absence of the GFP signal. Alternatively, it might be due to leaky expression of the postsynaptic reporter mtdTomato itself.

In view of the faint background noise that we observed in some brain regions, we decided to examine whether *retro*-Tango can be used in one of these regions, the central complex. The central complex is a series of interconnected neuropil structures that are thought to act as the major navigation center of the fly brain. The flow of information through the central complex indicates that it dynamically integrates various sensory cues with the animal’s internal state for goal-directed locomotion (Hulse et al., 2021). In the central complex circuitry, ellipsoid body-protocerebral bridge-gall (EPG) neurons have dendrites in the ellipsoid body (EB) and axons in the protocerebral bridge as well as in the lateral accessory lobes. EPGs are the postsynaptic targets of the ring neurons of the EB. They also form reciprocal connections with protocerebral bridge-ellipsoid body-noduli (PEN) neurons, protocerebral bridge-fan shaped body-round body (PFR) neurons and Δ7 interneurons(Hulse et al., 2021; Seelig & Jayaraman, 2013; Sun et al., 2017). When we initiated *retro*-Tango from the EPGs, we observed presynaptic signal in the predicted presynaptic partners (Figure 2c). Moreover, this signal was much stronger than the noise we observed in the absence of a driver, indicating that *retro*-Tango can indeed be used in brain regions with background noise. Further, the absence of labelling in any unexpected neuronal processes near the EPG cell bodies suggests that *retro*-Tango does not lead to false positive signal due to the presence of its ligand in neuronal somata (Figure 2–figure supplement 1). Finally, we do not observe presynaptic signal in starter neurons, indicating that expression of the *retro*-Tango ligand in a starter neuron does not activate the signaling pathway in the same cell (Figure 2–figure supplement 1).

We next sought to test the age-dependence of the presynaptic signal in *retro*-Tango. We initiated *retro*-Tango from the EPGs and examined the signal in adults at days 5, 10, 15, and 20 post-eclosion (Figure 3). We noticed that the signal accumulates and reaches saturation around day 10 post-eclosion. However, a similar analysis with GFs as the starter neurons indicated that the *retro*-Tango signal saturates later, around day 15 (Figure 3–figure supplement 1). Therefore, we concluded that the accumulation of the *retro*-Tango signal depends on the circuit of interest, and possibly, on the strength of the driver line being used. To be prudent, we examined adult flies 15 days post-eclosion for the remainder of the study.

**Figure 3.**
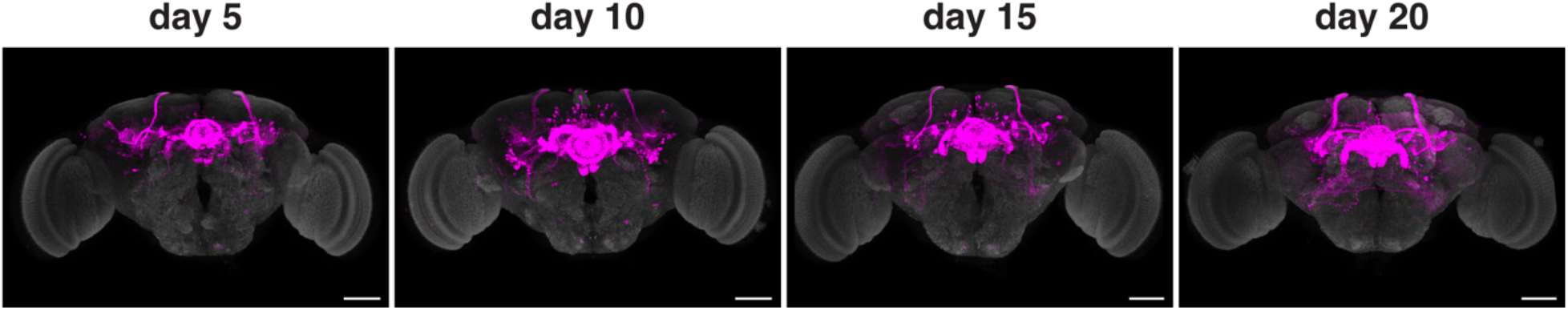
Age dependence of *retro*-Tango. The *retro*-Tango signal is observed in 5-day intervals upon ligand expression in the EPGs. The signal accumulates with time and saturates around day 10 post-eclosion. Presynaptic mtdTomato (magenta) and neuropil (grey). Scale bars, 50μm.

### Comparison of *retro*-Tango with the EM reconstruction of the female hemibrain

Having established the system in the GF and EPG circuits, we wished to benchmark it by comparing the presynaptic signal of *retro*-Tango with the EM reconstruction of the female hemibrain. In the connectome, we found 1101 neurons to be presynaptic to the giant fiber (Figure 4–figure supplement 1a). We observed fewer presynaptic neurons with *retro*-Tango (Figure 2a). Based on the EM reconstruction, the number of synapses that these 1101 neurons form with the GF ranges from 1 to 380. We, therefore, reasoned that the number of synapses that a given presynaptic neuron forms with the starter neuron affects whether it is labelled by *retro*-Tango. In other words, there is a threshold in the number of synapses that a presynaptic neuron makes with a starter neuron under which it cannot be labelled with *retro*-Tango. Neurons with fewer synapses than this threshold likely constitute the false negatives of *retro*-Tango. This threshold is probably affected by the circuit of interest and by the strength of the driver line.

To determine this threshold, we decided to count the presynaptic neurons of the GF revealed by *retro*-Tango using a nuclear reporter. In these experiments, we counted the neurons in each half of the brain focusing on the area that is covered by the connectome (Figure 4–figure supplement 1b, c). We counted five experimental GF *retro*-Tango brains and observed an average of 191(±42) neurons in this area. In six control brains from flies not carrying Gal4, we counted an average of 26(±11) neurons. We concluded that in this area, *retro*-Tango correctly labels approximately 165 neurons when initiated from the GF. Of the 1101 neurons that the connectome reveals as presynaptic to the GF, 341 have cell bodies in the area covered by the EM reconstruction. Therefore, *retro*-Tango identifies approximately half of these neurons. We analyzed the connectome data for these 341 neurons and found that 168 of them have each 17 synapses or more with the GF. Given that *retro*-Tango reveals approximately 165 neurons, we concluded that the threshold for *retro*-Tango to identify the presynaptic partners of the GF is 17 synapses (Figure 4–figure supplement 1a).

We subsequently used this newly determined threshold to sort the 1101 neurons revealed by the connectome as presynaptic to the GF and identified 265 neurons. We then plotted the skeletonizations of the EM segmentations of these 265 neurons (Figure 4a). When we initiated *retro*-Tango from the GF in females, we revealed a strikingly similar pattern (Figure 4b). It is noteworthy that we observe some differences in the *retro*-Tango signal between males and females. Based on the connectome, LPLC2s form an average of 13 synapses per neuron with the giant fiber (Ache et al., 2019). This is below the threshold, and indeed, we do not observe LPLC2s in females with *retro*-Tango (Figure 4b). By contrast, we do observe them in males (Figure 2a). This discrepancy could be explained by the location of the presynaptic mtdTomato reporter on the X-chromosome. Accordingly, the reporter expression level in males is higher compared to heterozygous females due to X-chromosome upregulation for dosage compensation(Gorchakov et al., 2009). Thus, the threshold in hemizygous males is significantly lower than in heterozygous females.

**Figure 4.**
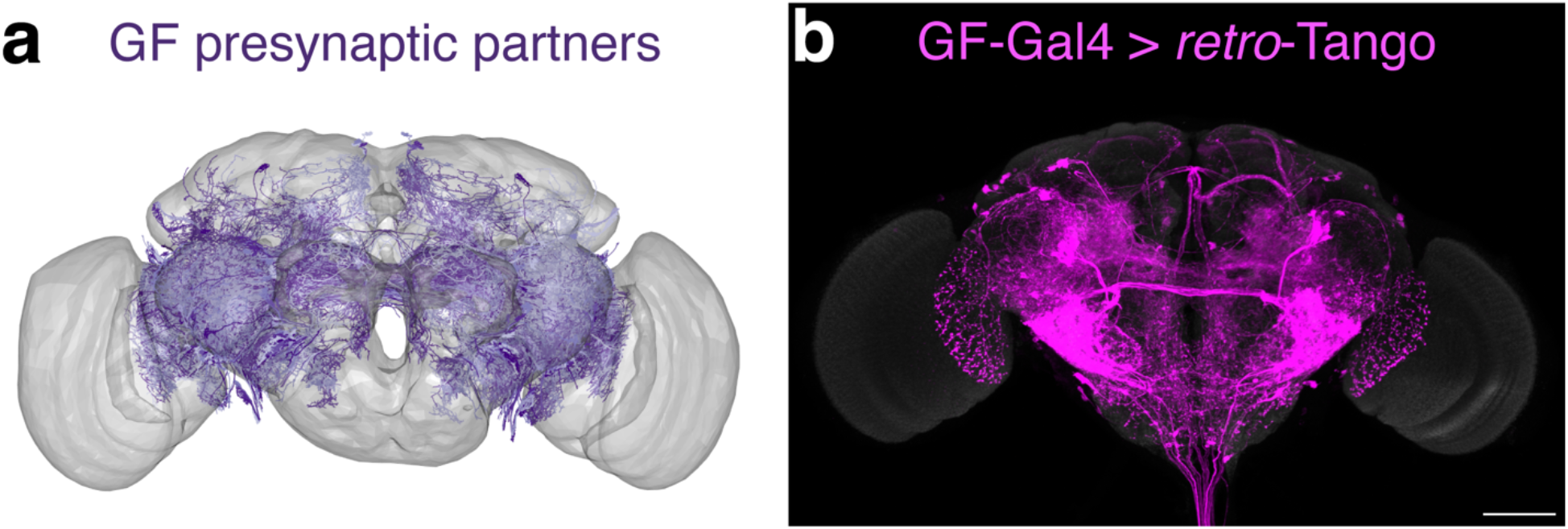
Comparison of the *retro*-Tango signal with the EM reconstruction of the female hemibrain. **(a)** Plotting of the skeletonizations of the EM segmentations of presynaptic partners that connect with the GF via 17 synapses or more. **(b)** Presynaptic partners of the GFs in a female fly as revealed by *retro*-Tango. Presynaptic mtdTomato (magenta) and neuropil (grey). Scale bar, 50μm. Note the high similarity between the patterns in both panels.

### Specificity of *retro*-Tango

Having benchmarked *retro*-Tango in tracing various connections, we sought to determine its specificity and reasoned that sexually dimorphic circuits would be apposite for this analysis. One such circuit involves the anterior dorsal neurons (aDNs), a pair of neurons in each hemisphere that receive inputs from distinct sensory systems in the two sexes. In males, the aDNs receive visual input, whereas in females, the input instead comes from the olfactory and thermo/hygrosensory systems (Nojima et al., 2021). Thus, we decided to use the sexual dimorphism in the inputs to aDNs for testing the specificity of *retro*-Tango. When we initiated *retro*-Tango from aDNs in males, we observed strong presynaptic signal in the central brain, and more importantly, in the visual system (Figure 5a). However, we did not observe presynaptic signal in LC10 neurons as would be predicted from this study (Nojima et al., 2021). A possible explanation for the absence of labeling in LC10s could be that the strength of connections between LC10s and aDNs is below the detection threshold of *retro*-Tango. Alternatively, LC10s may not be directly presynaptic to aDNs as the connections between these neurons were revealed by a non-synaptic version of GRASP (Gordon & Scott, 2009; Nojima et al., 2021). By contrast, in females, we observed two neurons in the lateral antennal lobe tracts, few neurons in the lateral horns (LHs), and neuronal processes in the suboesophageal zone (SEZ) as previously reported (Figure 5b). However, the signal in females is low, likely because they are heterozygous for the presynaptic reporter. Indeed, it seems that *retro*-Tango does not identify all the presynaptic neurons reported in females (Nojima et al., 2021). Nonetheless, the difference in the signal pattern between male and female brains demonstrates the specificity of *retro*-Tango.

**Figure 5.**
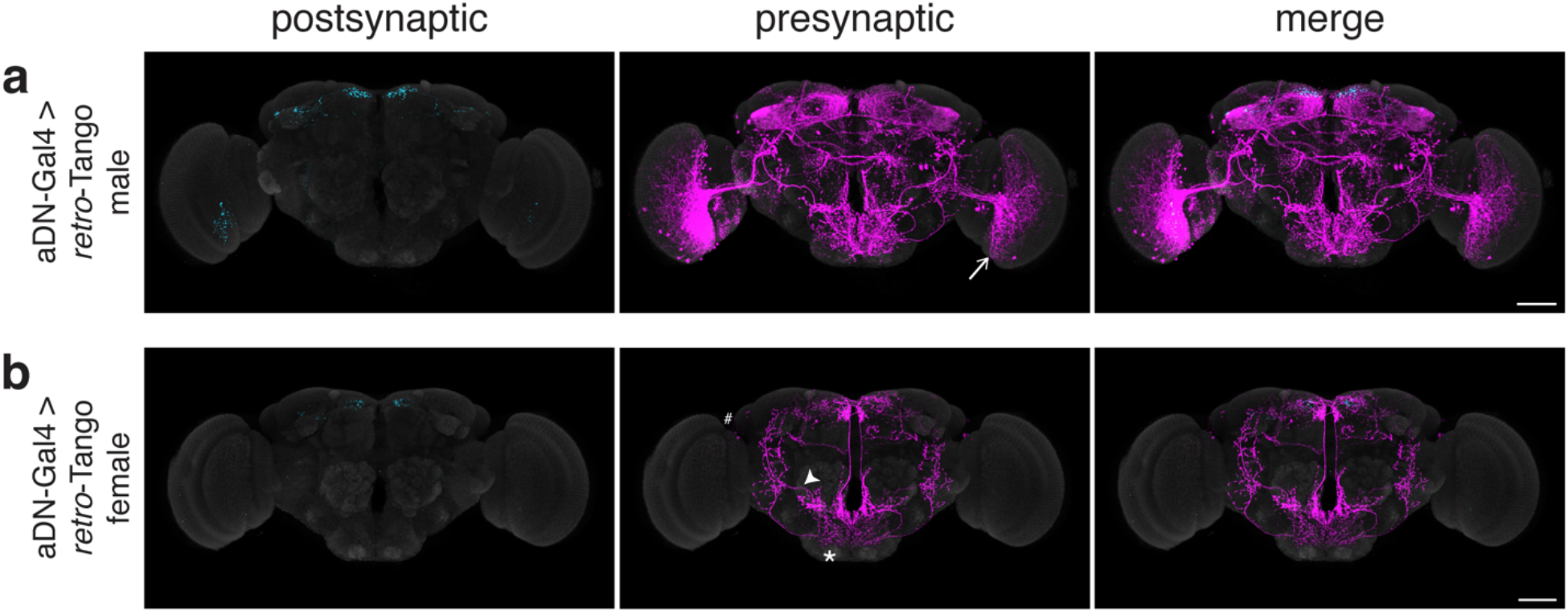
Revealing the specificity of *retro*-Tango in a sexually dimorphic circuit. **(a)** Initiating *retro*-Tango in aDNs in male flies reveals visual projection neurons (arrow) as presynaptic partners. **(b)** Initiating *retro*-Tango in aDNs in females results in presynaptic reporter expression in the lateral antennal lobe tract (arrowhead), the SEZ (asterisk), and the LH (hash). Postsynaptic GFP (cyan), presynaptic mtdTomato (magenta) and neuropil (grey). Scale bars, 50μm.

### Using *retro*-Tango to trace connections between the CNS and the periphery

Our experiments in the giant fiber, the central complex circuits and the aDNs established *retro*-Tango for tracing connections within the CNS. Next, we wished to examine whether *retro*-Tango can be used to trace connections between the CNS and the periphery. To achieve this, we turned to two well-characterized circuits: the sex peptide (SP) circuit and the olfactory circuit.

The SP circuit mediates the response of females to the presence of SP in the seminal fluid upon mating. SP is detected by the SP sensory neurons (SPSNs) located in the lower reproductive tract of females (Yapici et al., 2008). SPSNs project to the SP abdominal ganglion (SAG) neurons in the CNS to initiate the post-mating switch, a set of programs that alter the internal state of the female (Feng et al., 2014). Accordingly, initiating *retro*-Tango from SAG neurons reveals presynaptic signal in a pair of neurons in the lower reproductive tract, consistent with SPSNs (Figure 6a). This result confirms that *retro*-Tango can be used to reveal connections between the CNS and the periphery.

**Figure 6.**
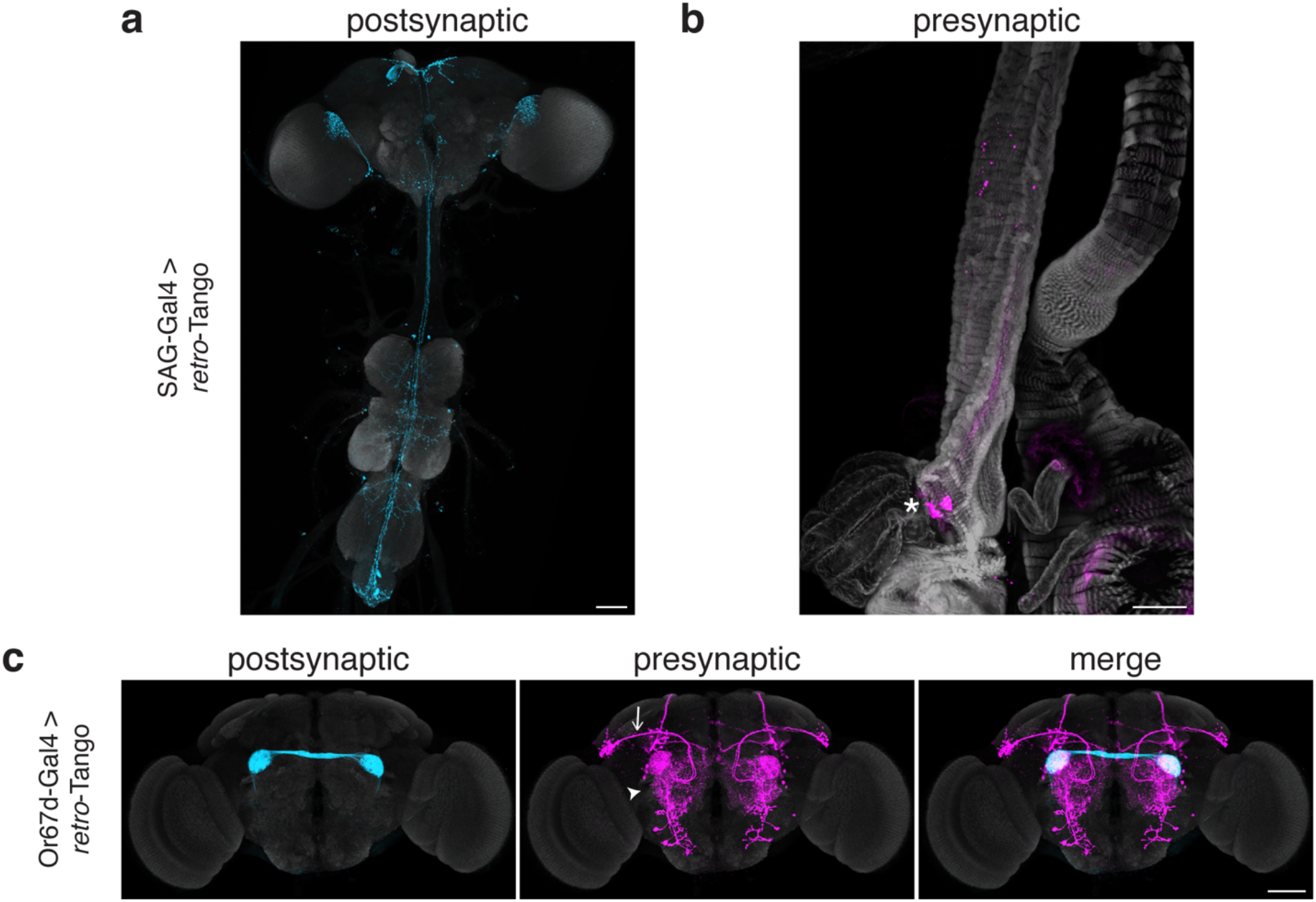
Tracing connections between the periphery and the CNS with *retro*-Tango. **(a)** Expression of the *retro*-Tango ligand in SAG neurons reveals **(b)** SPSNs (asterisk) as presynaptic partners. **(c)** When *retro*-Tango is initiated from Or67d-expressing ORNs, PNs (arrow) and LNs (arrowhead) are revealed as their presynaptic partners. Postsynaptic GFP (cyan), presynaptic mtdTomato (magenta) and neuropil **(a, c)**, or phalloidin **(b)** (grey). Scale bars, 50μm.

In the olfactory circuit, olfactory receptor neurons (ORNs) located in the antennae and the maxillary palps, the two olfactory sensory organs, project their axons to the antennal lobe, a brain region consisting of multiple neuropil structures called glomeruli. The ORNs that express the same olfactory receptor converge on the same glomerulus where they form synapses with lateral interneurons (LNs) and olfactory projection neurons (PNs). The PNs, in turn, relay the information to higher brain areas, primarily the mushroom body (MB) and the LH. Thus, in a simplistic model, the flow of sensory information is from the ORNs to the PNs while LNs form synapses with both neuronal types. However, all three neuronal types are interconnected via reciprocal synapses (Horne et al., 2018). Therefore, in this circuit, if we initiate *retro*-Tango in the ORNs, we expect to see presynaptic signal in the PNs and LNs. We, hence, sought to test *retro*-Tango in these reciprocal synapses. To this end, we initiated *retro*-Tango from a subset of ORNs that express the olfactory receptor Or67d and project to the DA1 glomeruli. We, indeed, observed presynaptic signal in PNs and LNs (Figure 6b). Together, these results confirm that *retro*-Tango can be used to reveal synaptic connections between the CNS and the periphery irrespective of the direction of information flow.

## DISCUSSION

In this study, we presented *retro*-Tango, a new method for retrograde transsynaptic tracing in *Drosophila. retro*-Tango is a versatile retrograde tracing method that can be used both as a hypothesis tester and a hypothesis generator. It shares many of its components with *trans-*Tango (Talay et al., 2017) and differs from it in the transmembrane protein with which the ligand is delivered. In *trans-*Tango a dNeurexin1-hICAM1 chimeric protein localizes the ligand to presynaptic sites such that it activates its receptor only in postsynaptic neurons across the synaptic cleft (Talay et al., 2017). By contrast, in *retro*-Tango the ligand is attached to mICAM5, a dendritic marker in *Drosophila* (Nicolai et al., 2010). Thus, driving the *retro*-Tango ligand in starter neurons activates the receptor in their presynaptic partners. This, in turn, triggers the signaling cascade culminating in reporter gene expression in the presynaptic neurons.

We used the GF circuit to validate *retro*-Tango since some of the known synaptic partners of the GFs can be easily identified. These experiments confirmed that *retro*-Tango correctly labels the expected presynaptic partners. In addition, we did not observe signal in the postsynaptic partners of the GFs, indicating that *retro*-Tango does not falsely label in an anterograde fashion. Further, driving ligand expression results in strong signal in the presynaptic neurons, while without a driver, the background noise is weak. We observed noise mainly in the MBs and the central complex with sporadic labelling in the VNC. To assess the utility of *retro*-Tango in these areas, we implemented it in the central complex. These experiments revealed presynaptic signal that can be easily discerned from the noise. That said, users should be cautious in drawing strong conclusions from *retro*-Tango experiments in these areas. As in *trans-*Tango (Talay et al., 2017), the panneuronal components are inserted at the attP40 docking site in the genome. It is noteworthy that the attP40 docking site that has recently been shown to cause problems in the nervous system, especially when homozygous (Duan et al., 2022; Groen et al., 2022; van der Graaf et al., 2022). Therefore, we advise against using the panneuronal components in a homozygous configuration. Likewise, users should be cautious when using Gal4 or split Gal4 lines inserted at the attP40 site.

The expression of mICAM5 is not entirely restricted to dendrites. Rather, it is also expressed in the somata, albeit at low levels (Nicolai et al., 2010). Hence, we were concerned that this would lead to labelling in neighboring neurons that are not true synaptic partners. However, our experiments in the central complex indicated that this is not the case. Nevertheless, caution should be taken especially when using strong drivers. It is also worth mentioning that we do not observe presynaptic labelling in the starter neurons, indicating that *retro*-Tango only works in trans.

Unlike *trans-*Tango(Talay et al., 2017), *retro*-Tango yields strong signal at 25ºC. This feature of *retro*-Tango is especially important as a recent study showed that the number of synaptic partners of a neuron and the number of connections with each partner are inversely correlated with rearing temperature (Kiral et al., 2021). Therefore, using *retro*-Tango at 25ºC prevents inconsistencies with other experiments run at this temperature. In addition, while like in *trans-*Tango (Talay et al., 2017) the signal in *retro*-Tango correlates with age, it accumulates faster. In some circuits, such as GF, the signal saturates at around day 15 post-eclosion, while in others, such as EPG, it only takes 10 days to saturate. The difference in saturation times could be due to the strength of the drivers or reflect the specific characteristics of the circuits. Therefore, users should determine the optimal age for analysis depending on the circuit studied and driver used. The availability of the annotated connectome data for the female hemibrain (Zheng et al., 2018) enabled us to benchmark the results obtained with *retro*-Tango and assess its sensitivity. To this end, we compared our results in the GF circuit to the annotated female hemibrain connectome (Zheng et al., 2018) (Figure 4). Our initial analysis indicated that *retro*-Tango falls short of revealing all the GF synaptic partners predicted by the connectome. Notably, some of these partners form single or few synapses with the GF. Therefore, it is possible that *retro*-Tango is not sensitive enough to reveal these weak connections. In our comparison, we determined the threshold for the number of synapses required for *retro*-Tango to correctly reveal a connection in the GF circuit. We applied this threshold to sort the presynaptic partners of the GF in the hemibrain connectome. When we plotted the neurons forming more synapses than the threshold, we observed a similar pattern to that revealed by *retro*-Tango.

One of the features that *retro*-Tango shares with *trans-*Tango is its modular design. In *retro*-Tango, this design provides genetic access to the presynaptic neurons. Therefore, the reporter can be readily swapped with an effector that allows for monitoring (Snell et al., 2022), activation, or inhibition of the presynaptic neurons. In addition, the modular design facilitates the adaptation of *retro*-Tango to other organisms. Notably, since using *retro*-Tango does not rely on a prior hypothesis regarding the identity of the presynaptic partners; it is flexible and general, and it can be used as a hypothesis generator. Presynaptic partners identified via *retro*-Tango can then be verified using orthogonal techniques. Therefore, *retro*-Tango is a significant addition to the toolkit for studying neural circuits that can open new avenues for circuit analyses.

## MATERIALS AND METHODS

### Fly Strains

Fly lines used in this study were maintained in humidity-controlled 25ºC incubators under standard 12h light/12h dark cycle. Flies were reared on standard cornmeal/agar/molasses media. Fly lines used in this study are: GF-split-Gal4 (von Reyn et al., 2014); *Or67d*^Gal4^ (Kurtovic et al., 2007); ss00090-Gal4 (Wolff & Rubin, 2018); SAG-split-Gal4 (ss51118) (Wang et al., 2021); aDN-split-Gal4 (Nojima et al., 2021); QUAS-nls-DsRed (Snell et al., 2022); QUAS-mtdTomato(3xHA) (this study); *retro-*Tango(panneuronal) (this study); *retro-*Tango(ligand) (this study).

### Generation of Transgenic Fly Lines

HiFi DNA Assembly (New England Biolabs #2621) was used to generate the plasmids used in this study. The plasmids were then incorporated into su(Hw)attP8, attP40 or attP2 loci using the ΦC31 system.

#### QUAS-mtdTomato(3xHA)

The QUAS-mtdTomato(3xHA) was amplified from UAS-myrGFP, QUAS-mtdTomato(3xHA) from the original *trans-*Tango study (Talay et al., 2017) using the following primers: cacggcgggcatgtcgacactagtgGTTTAAACCCAAGCTTGGATCCGGGTAATCGC and aactaggctagcggccggccttaattaaACTAGTGGATCTAAACGAGTTTTTAAGC. First, the plasmid pUASTattB (Bischof et al., 2007) was digested with SpeI and the whole mix was ligated in order to reverse the orientation of the attB site. The resultant plasmid was digested with BamHI and NheI and the PCR product was cloned into the plasmid via HiFi DNA Assembly. The final plasmid was incorporated into su(Hw)attP8.

#### *retro-*Tango(panneuronal)

The *retro-*Tango(panneuronal) plasmid was generated using the *trans-*Tango plasmid (Talay et al., 2017). The *trans-*Tango plasmid was digested with PmeI and AscI to remove the ligand and subsequently ligated to a dsDNA oligo mix containing AAACtaaGGCCGGCCcagGG and CGCGCCctgGGCCGGCCttaGTTT. The final plasmid was incorporated into attP40.

#### *retro-*Tango(ligand)

The *retro-*Tango(ligand) plasmid was generated using multiple components.

The 10xUAS to flexible linker sequence from the *trans-*Tango plasmid was amplified using ttgatttttttttttaagttggtaccCTCGAGCCTTAATTAACTGAAGTAAAG and cccagaaaggttcACTAGTATTCCCGTTACCATTG.

The mICAM5 sequence was amplified from fly lysates (Bloomington #33062 (Nicolai et al., 2010)) in two pieces using cgggaatactagtGAACCTTTCTGGGCGGACC & acagccatggaccGGCCACGCGCACTGTGAT and agtgcgcgtggccGGTCCATGGCTGTGGGTC & agttggtggcgccGGAAGATGTCAGCTGGATAGCGAAAACC.

The P2A sequence and the farnesylated GFP (GFPfar from addgene #73014) sequence was codon optimized and synthesized by ThermoFisher. It was, then, amplified using gctgacatcttccGGCGCCACCAACTTCTCC and ttattttaaaaacgattcatttaattaaTCAGGAGAGCACACACTTG primers.

The p10 sequence was amplified from the *trans-*Tango plasmid using tgtgctctcctgattaattaaATGAATCGTTTTTAAAATAACAAATCAATTGTTTTATAATATTCG TACG and acatcgtcgacactagtggatccggcgcgccGTTAACTCGAATCGCTATCCAAGC.

All five PCR products were then cloned into pUASTattB^11^ digested with BamHI and NheI. The final plasmid was incorporated into attP2.

### Immunohistochemistry, Imaging, and Image Processing

Dissection of adult brains, immunohistochemistry, and imaging were performed as described in the *trans-*Tango article (Talay et al., 2017) with modifications to accommodate for the clearing protocol. Unless otherwise stated adult male fly brains were dissected 15 days post-eclosion. Flies were cold anesthetized on ice and dissected in 0.05% PBST. Samples were fixed in 4%PFA/0.5% PBST for 30min, washed four times in 0.5% PBST, blocked in heat inactivated donkey serum (5% in 0.5% PBST) for 30min at room temperature. Samples were then treated with the primary antibody solution at 4ºC for two overnights. After four washes in 0.5% PBST at room temperature, samples were treated with secondary antibody solution at 4ºC for two overnights. After four washes in 0.5% PBST, samples were cleared following a previously published protocol(Aso et al., 2014). Reproductive system dissections were not subjected to the clearing protocol and were directly mounted on a slide (Fisherbrand Superfrost Plus, 12-550-15) using Fluoromount-G mounting medium (SouthernBiotech, 0100-01). Images were taken using confocal microscopy (Zeiss, LSM800) and were processed using the ZEN software from Zeiss. For nuclei counting, Imaris (version 9.1.2 Bitplane) was used. In all images, maximum projections are shown unless otherwise stated. The antibodies used in this study are as follows: anti-GFP chicken (Gift from Susan Brenner-Morton, Columbia University, 1:5,000), anti-RFP guinea pig (Gift from Susan Brenner-Morton, Columbia University, 1:10,000), anti-Brp mouse (nc82; DSHB; 1:50), donkey anti-chicken Alexa Fluor 488 (1:1000), donkey anti-guinea pig Alexa Fluor 555 (1:1000), donkey anti-mouse Alexa Fluor 647 (1:1000), Alexa Fluor 647 Phalloidin (Thermo Fisher A22287, 1:1000).

### Comparisons to the *Drosophila* Connectome

Data from the full adult fly brain (FAFB) electron microscopy (EM) volume (Zheng et al., 2018) was analyzed via the hemibrain connectome (Scheffer et al., 2020) using the natverse suite for neuroanatomical analyses in R (Bates, Manton, et al., 2020). The neuprintr package (Bates et al., 2022) was used to query the relevant cell types that we used as the starting populations for our *retro-*Tango experiments, as well as the identity of their presynaptic partners. Synaptic strength was determined as the total number of identified synaptic connections between the starting neuron and its presynaptic partner. Neurons in which the cell bodies were not traced as part of the hemibrain connectome were excluded from our counting experiments. To plot presynaptic cells, we used neuprintr to retrieve skeletonizations of their respective EM segmentations. Since the hemibrain connectome contains only segmentations of neurons from one side of the brain, we used natverse tools for bridging registrations to mirror the presynaptic neurons across the sagittal plane to the opposite hemisphere. Briefly, skeletonizations were translated from the FAFB space to the JFRC2 template (Jenett et al., 2012), which contains information for translating coordinates across sagittal hemispheres. Mirrored skeletonizations were then translated back to the FAFB space and plotted alongside the unmirrored data.

## Acknowledgments

We would like to acknowledge Dr. Cagney Coomer, Dr. Marnie Halpern, Dr. Jennifer Li, Dr. Karla Kaun, Daria Naumova, Dr. Drew Robson, and Dr. Rahul Trivedi for helpful discussions. We would like to thank Dr. Alexander Fleischmann and the members of the Barnea Laboratory for critical reading of the manuscript. We are grateful to Dr. Stephen Goodwin and Susan Morton for sharing reagents. This work was supported by NIH Brain Initiative grant NIH RF1MH123213 (G.B.), Brown University Carney Institute for Brain Science, Suna Kıraç Fund for Brain Science (D.S.), Brown University Carney Institute for Brain Science, Graduate Award in Brain Science (D.S.) and NIH/NIDCD award F31DC019540 (A.M.C.). Stocks obtained from the Bloomington *Drosophila* Stock Center (NIH P40OD018537) were used in this study.

## Author contributions

A.S., R.A.M., A.M.C., D.S., A.M.O. and G.B. conceptualized the study. A.S., M.T. and G.B. devised the methodology. A.S., R.A.M., A.M.C., D.S., and A.M.O. contributed to the investigation and visualization of the results. A.S. and G.B. administered the project. A.S., R.A.M. and G.B. wrote the manuscript. The funding was obtained by, and the project was supervised by G.B.

## Competing interests

Authors declare that they have no competing interests.

## Additional information

All new fly strains will be deposited to Bloomington *Drosophila* Stock Center. Correspondence and requests for materials should be addressed to G.B.

## Notes

### Competing Interest Statement

The authors have declared no competing interest.

