## Supplementary Figures for "*retro*-Tango enables versatile retrograde circuit tracing in *Drosophila*"

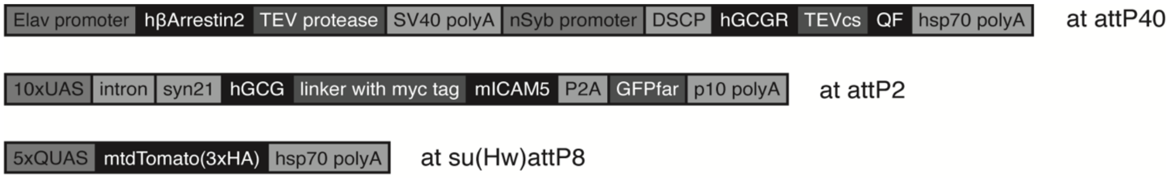


Figure 1–figure supplement 1 **The genetic components of *retro*-Tango**
Details of the genetic components used in *retro*-Tango are shown. Schematics are not drawn to scale. Elav: *Drosophila melanogaster* panneuronal promoter; polyA: polyadenylation signal; nSyb: *Drosophila melanogaster* panneuronal promoter; DSCP: Drosophila Synthetic Core Promoter; hGCGR: human Glucagon Receptor; TEVcs: cleavage site for N1a protease from the Tobacco Etch Virus; UAS: Upstream Activating Sequence for Gal4; hGCG: human Glucagon analogue with enhanced receptor binding; mICAM5: mouse intercellular adhesion molecule 5; P2A: 2A peptide from porcine teschovirus-1; GFPfar: farnesylated Green Fluorescent Protein; QUAS: Upstream Activating Sequence for QF.

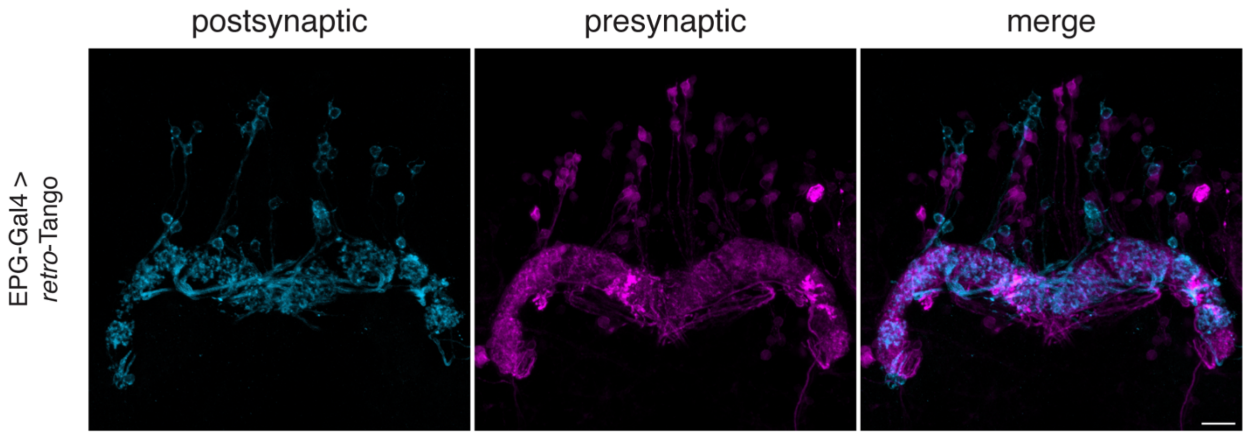


Figure 2–figure supplement 1 **No false positive signal is observed in neighbouring neurons with *retro*-Tango**

When *retro*-Tango is initiated from EPG neurons, the ligand present in the cell bodies does not lead to false positive presynaptic signal in neighbouring neurons. For clarity, only a subset of the z-stack projection is shown. Postsynaptic GFP (cyan), presynaptic mtdTomato (magenta). Scale bar, 10µm.

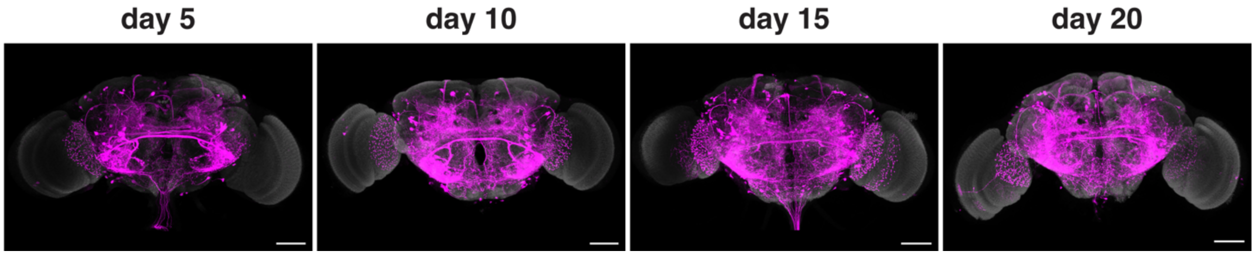


Figure 3–figure supplement 1 **Age dependence of the *retro*-Tango signal in the presynaptic partners of the GFs**
*retro*-Tango signal is observed in 5-day intervals upon ligand expression in the GFs. The signal saturates around day 15 post-eclosion, later than in the EPG circuit (Fig. 3). Presynaptic mtdTomato (magenta) and neuropil (grey). Scale bars, 50µm.

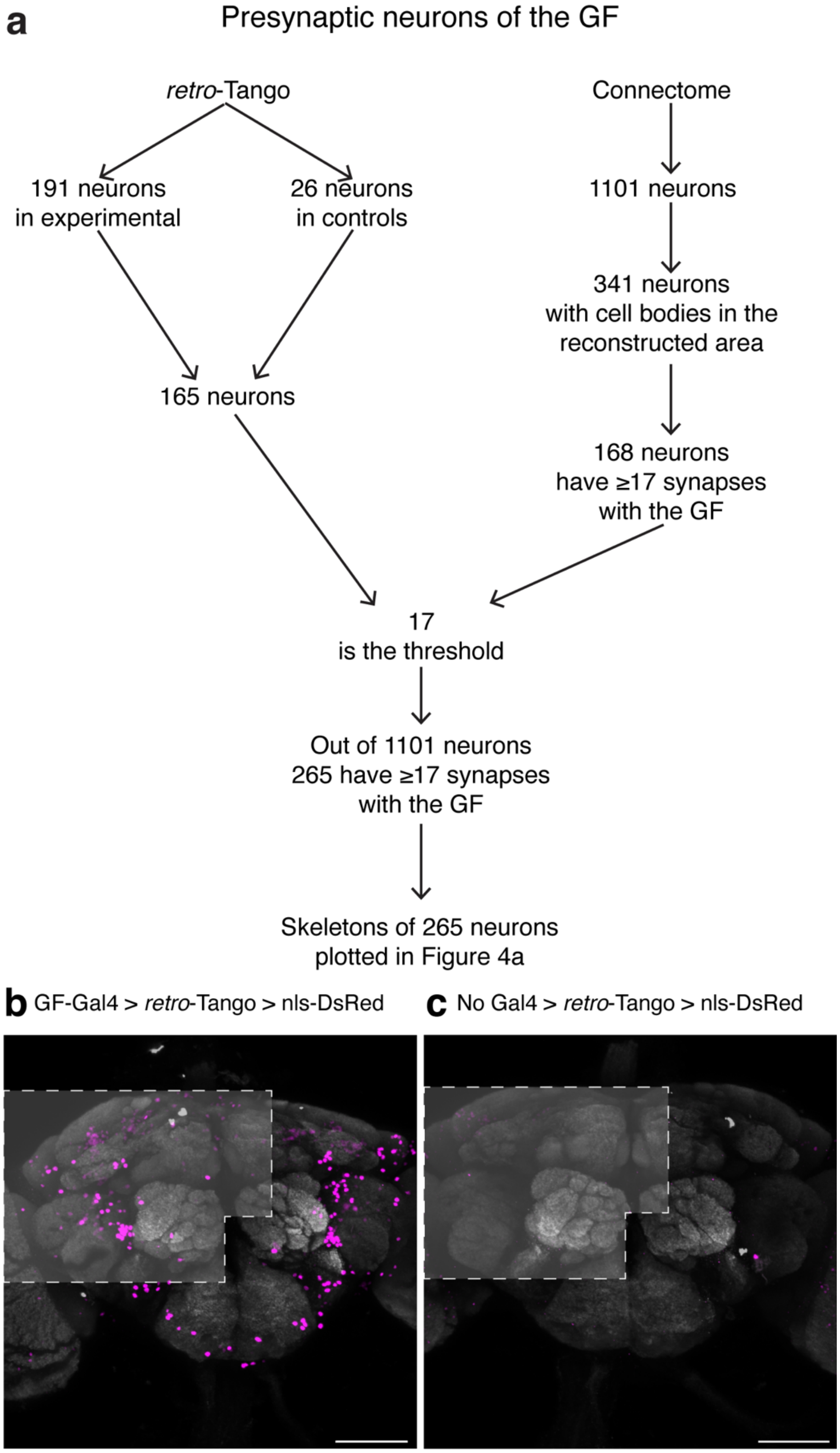
a
Presynaptic neurons of the GF
retro-Tango
Connectome
191 neurons in experimental
26 neurons in controls
1101 neurons
341 neurons with cell bodies in the reconstructed area
165 neurons
168 neurons have ≥17 synapses with the GF
17 is the threshold
Out of 1101 neurons 265 have ≥17 synapses with the GF
Skeletons of 265 neurons plotted in Figure 4a
b GF-Gal4 > retro-Tango > nls-DsRed
c No Gal4 > retro-Tango > nls-DsRed

Figure 4–figure supplement 1 **Methodology for the comparison of *retro*-Tango results with the hemibrain connectome**

**(a)** Flowchart explaining the steps in the comparison. **(b)** Driving *retro*-Tango from the GFs results in nuclear staining in an average of 191 neurons in ten hemibrains. **(c)** In the absence of a Gal4 driver, *retro*-Tango has background nuclear staining in 26 neurons. The areas analysed are marked in light grey based on the approximate regions covered by the published hemibrain connectome. Presynaptic mtdTomato (magenta) and neuropil (grey). Scale bars, 50µm.
